# Living in a metal-rich world: Enhanced growth and reduced metal accumulation in *Fusarium* fungi from the Kiirunavaara iron ore mine

**DOI:** 10.64898/2026.07.09.737466

**Authors:** P. Madsen, N. Hensen, M Orsucci, Hanna Johannesson

## Abstract

**Background:** Human activities such as mining generally lead to increased heavy metal concentrations in the environment. While traditional remediation techniques are often costly, the use of fungi as bioremediators, known as mycoremediation, is increasingly gaining attention as a sustainable approach for removal of heavy metals. Here, we evaluated heavy metal levels inside the Kiirunavaara iron ore mine in Northern Sweden and analysed fungal responses to various metal concentrations by comparing growth and metal uptake in mine-derived isolates and closely related control isolates.

**Results:** Sediments inside the mine were enriched in heavy metals compared to those from the outlet of the mine to natural lakes. Six *Fusarium* isolates were recovered from contaminated mining environments: five isolates from inside the mine were identified as *Fusarium oxysporum*, and one isolate from the outlet was identified as *Fusarium tricinctum*. Isolates from the mine and outlet showed overall higher survival and biomass production in presence of copper, iron, and zinc across a range of concentrations (up to 1000 mg/L) compared to control isolates. At the same time, these isolates often exhibited reduced relative metal uptake. As a result, mycoremediation potential, assessed as total uptake in the grown mycelium, was isolate-dependent.

**Conclusions:** Based on these results, we conclude that *Fusarium* isolates from the Kiirunavaara mine show increased growth in media enriched with heavy metals compared to closely related control isolates. We additionally show that mycoremediation potential is not necessarily associated with environmental origin. Instead, mycoremediation potential should be evaluated on a case-by-case basis for each isolate and based on specific needs for mycoremediation.

## Introduction

Heavy metals are among the most common contaminants affecting soil and groundwater in Europe (1). Some heavy metals, such as iron, copper and zinc, are essential and required at trace levels for cellular functioning, but can become toxic even at low concentrations (2). Heavy metals cannot be degraded through chemical or biological processes, allowing them to persist in the environment and accumulate in higher trophic levels of the food chain (3).

Anthropogenic activities, such as mining, can increase heavy metal concentrations to dangerous levels (4). Chemical methods to manage heavy metal contamination are generally costly and present several limitations, such as generating large volumes of sludges that require disposal (5). As a result, bioremediation methods that use plants, bacteria, or fungi to remove or mitigate the harmful effect of heavy metal contaminants are gaining increasing attention (5–9). Fungal-based bioremediation, also called mycoremediation, of heavy metals is particularly promising due to fungi’s ability to take-up and biotransform heavy metals (see e.g. (10)). Mycoremediation is considered a cost-effective process that does not leave harmful by-products, and has shown promising results (11). Multiple *Fusarium* species, for example, show resistance to heavy metals (10,12,13), making them suitable candidates to study mycoremediation potential.

In fungi, long-term exposure to heavy metal rich environments can influence tolerance to heavy metals through physiological and adaptive process (14). Heavy metal tolerance is usually evaluated by assessing growth responses across a gradient of metal concentrations relative to metal-free media (see e.g.(14)). However, mycoremediation potential depends not only on survival and growth under metal stress, but also on the organism’s ability to remove metals from the environment (15). Mycoremediation potential is normally researched only in fungi isolated from contaminated environments, as these are often assumed to be more resistant to high levels of heavy metals than those from non-contaminated sites (10–13). Tolerance and uptake are not always positively correlated, however. Some mushroom forming fungi are capable of growing in soils with elevated levels of heavy metals by maintaining homeostasis through limiting intracellular accumulation (16,17). Such negative relationships between heavy metal tolerance and uptake suggests that mycoremediation potential should be evaluated on a case-by-case basis, rather than only being inferred from highly tolerant strains isolated from contaminated environments. Despite these insights, few studies include assessment of both tolerance and uptake across closely related fungi from different environments.

The Kiirunaavara Iron Ore mine in Sweden is the largest underground iron-ore mine in the world, and has been pivotal for Sweden’s steel production since the 1950’s. The mine therefore represents a valuable system to study fungal responses to long-term exposure to heavy metals. This study investigates heavy metal tolerance and uptake in six *Fusarium* isolates recovered from sediments inside the mine and from its outlet. Their performance was compared to four closely related control isolates to evaluate the interplay between environmental origin, species identity, tolerance, and uptake in shaping mycoremediation potential. Specifically, we assessed survival, biomass, and relative and absolute metal uptake in response to concentrations of copper (Cu), iron (Fe) and Zinc (Zn) up to 1000 mg/L. We hypothesized that isolates from mining environment would exhibit a higher tolerance to heavy metal stress, but that increased tolerance would not necessarily translate into a better mycoremediation potential, i.e., the ability to uptake heavy metals in the growing mycelium.

## Methods

### Study area

The Kiruna mine Iron Ore mine in Sweden is operated by Luossavaara-Kiirunavaara AB. The mine is structured vertically, with industrial activity following the iron ore body (Figure 1). The industrial runoff from the mining activity consists of water loaded with salts, sand, and particle bound elements, and is continuously pumped in a sequential pumping chain. The water is pumped with a maximum flow of ∼25m^3^/minute, from the bottom pumping pit towards the top of the mine. There are in total five pumping pits underground, which accumulate the runoff and are periodically emptied when full. Sediments are taken to landfills and the runoff is led to surface level and into a sedimentation basin, followed by a clarification basin, before being let out into a stream (the outlet) which leads to a natural lake. For this study, heavy metal levels were assessed in sediment samples taken from pumping pits KP90, KP112, KP136, KP153, and from the outlet (Figure 1), and analysed with ICP-SFMS according SS-EN ISO 17294-2:2016 (18). Due to difficulty of access at time of sampling, sediment samples from KP57 were not included in the assessment. Reference levels of heavy metal concentrations for freshwater sediments were obtained from the Geological Survey of Sweden (obtained March 2026) and are given in Table 1 (19). Lake Jutsajaure, located in Gällivare municipality, was used as a reference lake not directly affected by anthropogenic activities (19). Table 1 also lists the average estimates of lakes in Norrbotten County, which is the northern region in Sweden where the mine is located, and the national average (Figure 1, Table 1).

**Figure 1:**
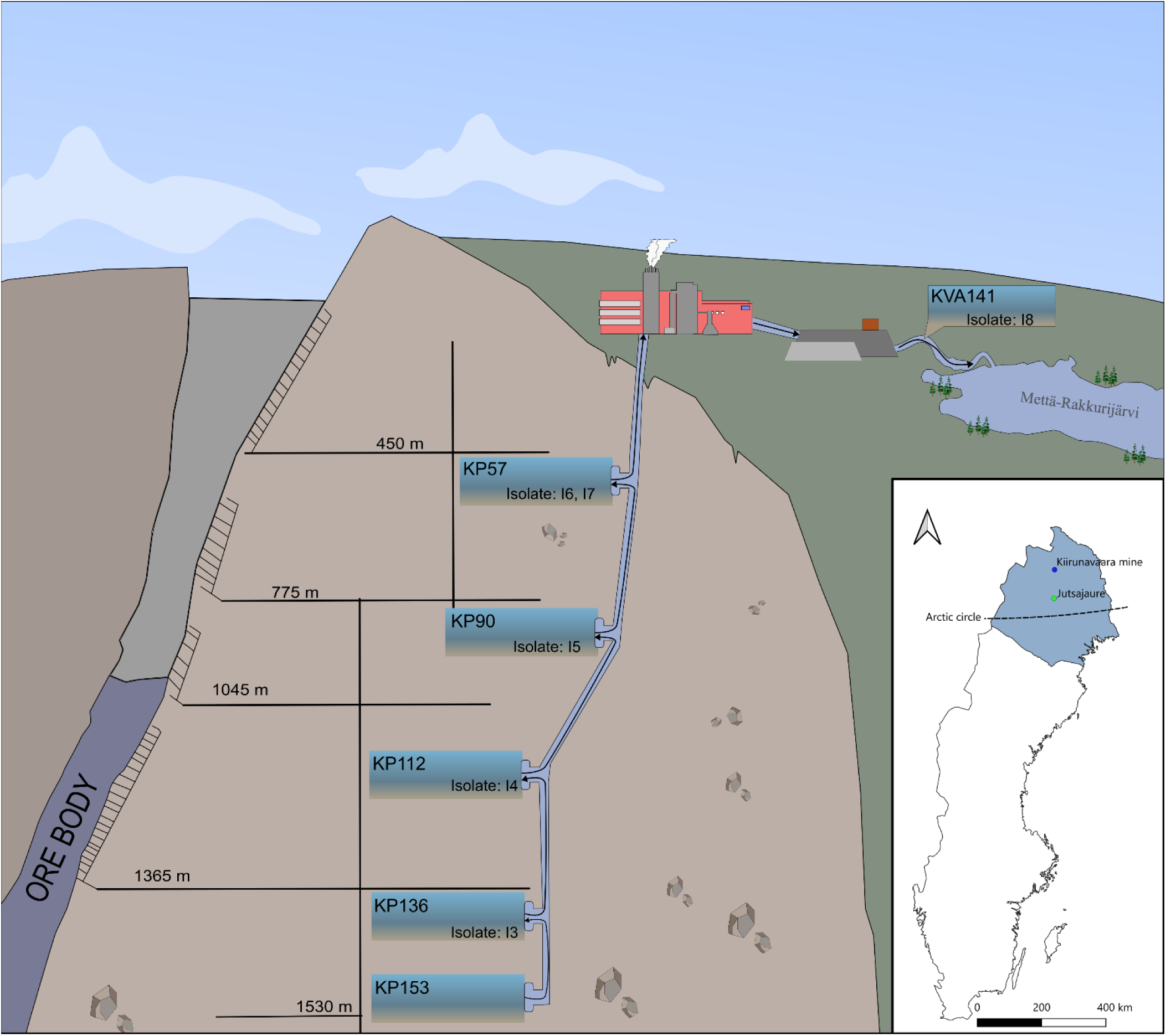
Graphical overview of the structure of the Kiruna Iron Ore mine and geographical location. The pumping pits (KP57, −90, −112, −136 and −153) and outlet (KVA141) are shown. The depth of the pumping pits is approximate, and fungal isolates are from the various pumping pits in the mine and the outlet. Processing water transport runoff and sludge to the pumping pits, flow of water indicated by arrows. Upon saturation of processing water, the water is pumped up to surface level and led through multiple cleaning steps before being released into nearby natural lakes. The map shows Norrbotten county highlighted in blue, the position of the arctic circle, the Kiirunavaara mine (67.843504004N, 20.167978015E) and the reference lake Jutsajaure (67.050756988N, 19.957315795E) are indicated.

**Table 1:** Average concentrations of heavy metals inside the mine, at the mine’s outlet, in Swedish reference lake Jutsajaure, the average for Norrbotten county, and the National average of freshwater lakes.

| Dry substance<br>(mg/kg) | Mine (n =4)<br>Mean ± std dev | Outlet (n = 1) | Jutsajaure<br>Mean (n = 1) | Norrbotten<br>Mean (n = 306) | National<br>Mean (n = 2024) |
| --- | --- | --- | --- | --- | --- |
| As | 4.8 ± 1.0 | 7.1 | 4.53 | 95.4 | 12.9 |
| Cd | 0.1 * | < 0.05* | 0.46 | 3.7 | 1.6 |
| Co | 67.4 ± 2.3 | 26.4 | 9.1 | - | 20.6 |
| Cr | 10.8 ± 5.6 | 52.2 | 14.0 | 29.2 | 52.1 |
| Cu | 166.5 ± 39 | 57.5 | 14.0 | 30.9 | 77.9 |
| Hg | <0.05 * | < 0.05* | 0.1 | 0.2 | 0.2 |
| Mn | 833.3 ± 330.4 | 660.0 | 478.0 | 10071.0 | 1124.0 |
| Ni | 113.0 ± 18.4 | 34.4 | 9.87 | 23.9 | 35.7 |
| Pb | 6.8 ± 2.6 | 5.4 | 19.2 | 33.6 | 58.0 |
| V | 460.8 ± 157.8 | 140.0 | 25.8 | - | 54.3 |
| Zn | 250.8 ± 152.6 | 37.4 | 129.0 | 150.0 | 428.0 |
| Fe | 226750.0 ± 62590.6 | 57500.0 | 62337.0 | 50728.0 | 43804.4 |

\* measurements below detection level. Reported level is the detection level.

### Isolation and identification of sediment fungi

To obtain fungal isolates from the mine, sediment samples were taken from all five pumping pits, as well as the outlet (Figure 1). These sediment samples were mixed with liquid mineral salt medium (NaCl (0.5 g/L), (NH4)2SO4 (0.1 g/L), NaNO3 (0.2 g/L), MgSO4 7H2O (0.025 g/L), K2HPO4 3H2O (1.0 g/L), and KH2PO4 (0.4 g/L), as listed in (20), supplemented with synthetic engine oil (1% v/v). From the resulting cultures, aliquots of 20 µL were taken and plated onto 2% Potato Dextrose Agar (PDA) plates and colonies of different fungi and bacteria were allowed to grow. Based on macroscopic characteristics, 14 morphologically distinct fungal isolates were selected for further investigation, and these were transferred individually to new 2% PDA-plates. To establish pure cultures, a combination of mechanical cleaning and antibiotic treatment was employed, as described by Shi et al. (21). In short, a single fungal hyphal tip was transferred from each of the cultures to a central well of 2% PDA plates with 0.1% of penicillin-streptomycin, allowing fungal hyphae to grow outward from the central well for isolation. This cleaning process was repeated up to six times for each culture.

For molecular identification of fungal isolates, and to verify absence of bacterial contaminants, we used a PCR-based approach. See supplementary methods for details on DNA extraction, PCR analysis and sequencing methods. Fungal cultures were assumed to be free of bacterial contamination based on the absence of bacterial PCR products at the start of the experiment.

### Fungal identification, selection of control isolates, and phylogenetic reconstruction

Taxon identity of the fungal isolates obtained from the mine was inferred by comparing ITS sequences to the NCBI database using BLASTN (22). Six isolates were assigned to *Fusarium* spp. and formed the basis for further investigation. Based on the ITS1 region, five were identified as *Fusarium oxysporum*, and one was determined to be *Fusarium tricinctum.* To identify suitable controls from non-industrial environments, the six *Fusarium* ITS sequences were blasted against the filamentous fungal CBS ITS barcode data set of Vu et al. (2019) (23). For *F. oxysporum*, the top three blast hits of available CBS strains were selected as controls. For *F. tricinctum,* we compared our ITS sequence to ITS sequences of *F. tricinctum* CBS sequences from the NCBI nucleotide database to identify the best control isolate. Three of the four controls were isolated from plants, and the fourth from unknown substrate. Pure cultures of the controls were obtained from the Westerdijk institute (Table S1).

To investigate the phylogenetic relationship between the controls and the fungi isolated from the mining environment, we performed phylogenetic analyses of the ITS1 sequences, with *Microascus trigonosporus* (CBS 218.31) as the outgroup. ITS sequences were aligned with MAFFT V7.525, with automatic model selection (“—auto”) (24). The alignment was visually examined with Jalview V2 (25), showing large similarity between ITS sequences and no need for manual curation or further trimming. Maximum likelihood analysis was performed in R v4.6.0 with the ape and phangorn packages (v5.8-1 and v2.12.1; (26–29)). Automatic model selection revealed TPM2+G4 as best fitting model. The support for each internal branch was evaluated with 1000 bootstrap replicates.

### Fungal growth assays

The four control isolates and six mine isolates were maintained on 4% malt extract agar at ∼23°C. Growth was assessed for the ten isolates at concentrations of 200 mg/L, 600 mg/L and 1000mg/L for copper, zinc and iron. Fungal isolates were inoculated in 100 mL 4% malt extract broth (MEB), enriched with filter sterilized stock solutions (pore size 0.2 µm) of copper (CuSO4·5H2O), zinc (ZnSO4·7H2O), or iron (FeSO4·7H2O) to reach the desired concentrations. MEB without metal salts and uninoculated cultures were used as control media.

The pH of the media (n=3 for all measurements) was assessed across various metal concentrations without fungal inoculates. The lowest mean pH, 3.35, occurred at Cu 1000 mg/l, while the highest, 5.46, was observed in the MEB media without metal supplementation. For zinc, the average pH values were 4.94 at both 200 and 600 mg/l, and 4.86 at 1000 mg/l. Copper exhibited a decreasing trend, with average pHs of 4.05, 3.51, and 3.35 at 200, 600, and 1000 mg/l, respectively. Iron’s pH values were 5.13, 4.94, and 4.80 at the same concentrations.

For inoculation, a small agar plug (roughly 3 mm in diameter) containing mycelium from the edge of a growing colony was transferred to the culture flasks containing liquid medium. Cultures were incubated on a shaker at 60 rpm at room temperature (∼23°C) for nine days. All growth tests were conducted in triplicate, yielding three biological replicates per isolate and growth condition.

Following incubation, cultures were vacuum filtered using grade 3 cotton linter filters (particle retention >10 µm), and mycelium was washed with 50 mL of deionized water. Fresh weight of the mycelium was obtained by weighing precipitate for all isolates with fungal growth. All cultures that showed growth were used for further analysis of dry weight by ALS Scandinavia. Dry weight was obtained by freeze-drying fresh mycelium using TS-105 according to SS-EN 15934:2012 (18) 2024). There was a high, significant correlation between fresh and dry weight, as assessed with Spearman’s rank correlation (ρ = 0.72, *p < 0.01*; function cor.test, package stats), supporting the use of dry weight for further analyses.

### Trait analysis and statistical analysis

Four traits were measured: survival (growth versus no growth), biomass (dry weight of mycelium), relative uptake (mg heavy metal per kg dry weight mycelium), and absolute uptake (mg heavy metal in grown mycelium). All analyses were performed in R (v4.4.1) using RStudio (28,29). Because fungal species and isolate origin were not fully crossed in the experimental design, they were combined into a single factor (“group”) with four levels: *F. oxysporum*-control, *F. tricinctum*-control, *F. oxysporum*-mine, and *F. tricinctum*-outlet.

Survival was analysed using generalized linear mixed models (GLMMs) with binomial error distribution and logit link, fitted with the glmmTMB package (30). Fixed effects included metal type, metal concentration (continuous variable, centered and scaled), and group. Fungal isolate was included as a random effect. Because of convergence issues due to a quasi-complete separation in the full interaction model (Table S3), reduced models including two-way interactions were fitted, and the most parsimonious model was retained using likelihood ratio tests.

Biomass was analysed using linear mixed-effects models (LMMs) fitted with the lme4 package (31). Dry weight was log-transformed (log1p) and explained by group, metal, scaled concentration, and their interactions as fixed effects, with fungal isolate as a random effect. Additional analyses were performed on samples with growth (weight > 0) to assess whether patterns of biomass response to metal concentration were consistent when excluding non-growing samples. For all models, significance of fixed effects was assessed using Type II Wald chi-square tests with the Anova function (car package; (32)). Posthoc comparisons and estimation of concentration-response slopes were performed (emmeans package; (33)).

Relative and absolute uptake were assessed for samples with sufficient growth (> 3.5g fresh mycelial weight; Table S1-S2). Due to limited sample size, these traits were not statistically analysed but described qualitatively. Relative uptake was measured by ALS Scandinavia using ICP-SFMS and expressed as mg of metal per kg of dry mycelial weight, with a minimum of 3.5g of biomass needed for analysis. Absolute uptake was determined by multiplying the relative uptake by the dry weight of the mycelium.

## Results

Analysis of sediments from the mine’s pumping pits revealed elevated levels of multiple heavy metals when compared to concentrations assessed at the mine’s outlet, including high mean concentrations of the essential heavy metals copper, iron and zinc (Table 1). All metals except arsenic and chromium showed higher concentrations in the mine pumping pits compared to the outlet. Iron concentrations were particularly high, with average levels of 226750 mg/kg in the sediments of pumping pits, and 57500 mg/kg in the outlet sediment. In comparison, the average level of iron in the sediments from the reference lake Jutsajaure was 62337mg/kg and the National average was 43804.4 mg/kg (Table 1). The mine showed higher concentrations of iron, copper and zinc in the mine than in the reference lake Jutsajaure and the Norrbotten county average. In addition, the mine contained higher values of heavy metals such as Ni, V and Co in comparison to the reference lake Jutsajaure, Norrbotten county, and the National average (Table 1).

### Phylogenetic relationship of mine and control isolates

Maximum Likelihood analysis of the ITS1 sequences of six *Fusarium* isolates from the mine and the four closely related controls produced a tree with two clades separating the two taxa, *Fusarium tricinctum* and *F. oxysporum* (Figure 2). In the *F. oxysporum*-clade, which included both mine and control isolates, these did not cluster separately, suggesting that observed differences in growth and uptake between the groups are unlikely to be explained solely by phylogenetic relatedness.

**Figure 2:**
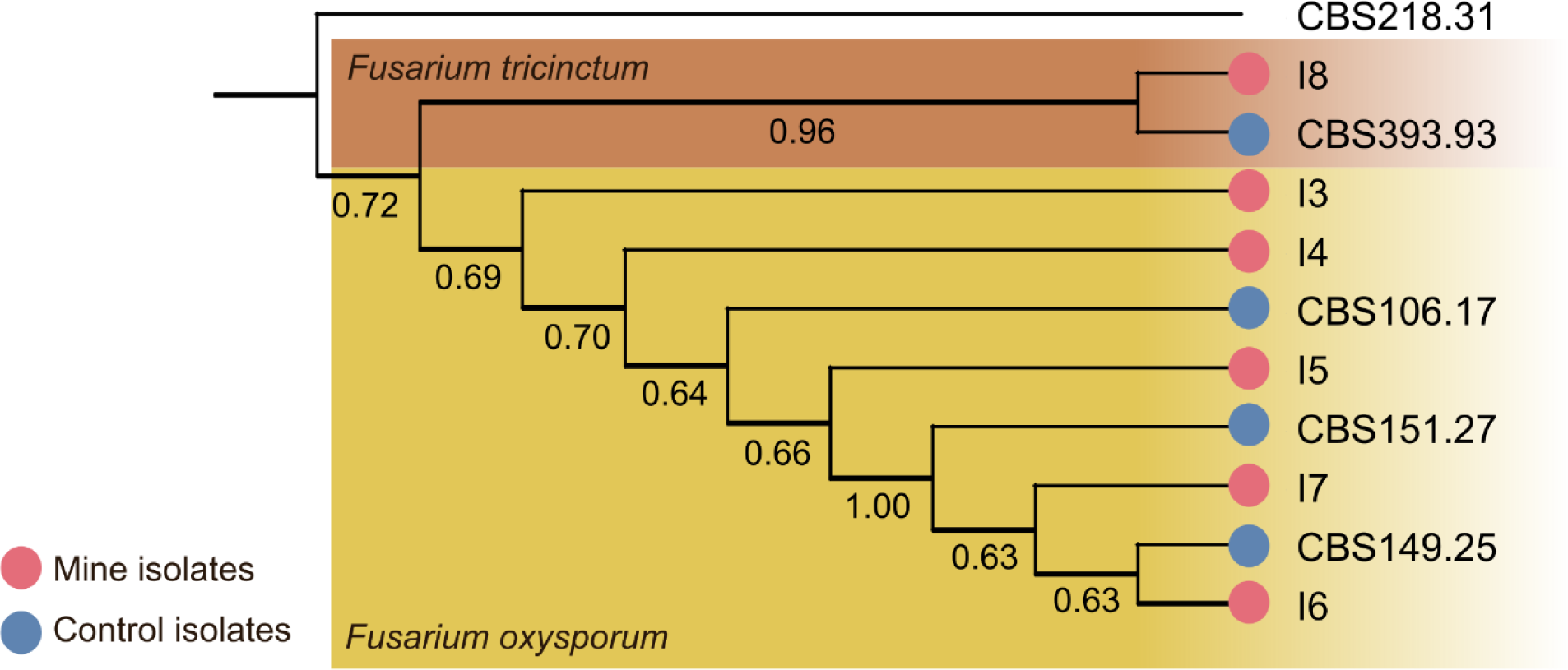
ITS tree of assessed fungal isolates from the Kiirunavaara mine and controls. Branches show bootstrap values evaluated with 1000 bootstrap replicates. Isolates from the mining environment are indicated with red circles on the branch tips and Isolate (I) number, while those from control environments are indicated with blue circles on the branch tips and by CBS (CBS) number.

### Increased survival and biomass in several isolates derived from the mining environment

Survival probability was significantly affected by group, metal, and concentration (Table 2; Table S2-S4). The model including the metal × concentration interaction provided a better fit than an additive model (LRT, χ² = 6.47, df = 2, *p* = 0.039) and was retained. Within *F. oxysporum*, isolates from the mine showed higher survival than control isolates (Tukey-adjusted, *p* < 0.001), whereas no clear difference was observed for *F. tricinctum.* The effect of concentration on survival varied among metals, as indicated by a significant metal × concentration interaction (Table 2). Survival declined most strongly with increasing iron concentration, whereas the decrease was weaker in zinc (Fe vs Zn, *p* = 0.021). Copper showed an intermediate response (Cu vs Zn, p = 0.094; Cu vs Fe, *p* = 0.587).

**Table 2:** Effect of model parameters in different assessed models.

| **a. Survival model** |  |  |  |
| --- | --- | --- | --- |
| **Effect** | **df** | **Chisq** | **p-value** |
| Group | 3 | 33.45 | <0.001 |
| HM | 2 | 12.14 | 0.002 |
| Concentration | 1 | 30.11 | <0.001 |
| concentration × HM | 2 | 7.32 | 0.026 |
| **b. Biomass model (all data)** |  |  |  |
| **Effect** | **df** | **Chisq** | **p-value** |
| Group | 3 | 29.45 | <0.001 |
| Concentration | 1 | 301.53 | <0.001 |
| HM | 2 | 57.07 | <0.001 |
| group × concentration | 3 | 60.12 | <0.001 |
| group × HM | 6 | 40.52 | <0.001 |
| concentration × HM | 2 | 85.94 | <0.001 |
| group × concentration × HM | 6 | 35.08 | <0.001 |
| **c. Biomass model (among survivors)** |  |  |  |
| **Effect** | **df** | **Chisq** | **p-value** |
| Group | 3 | 17.94 | <0.001 |
| Concentration | 1 | 99.47 | <0.001 |
| HM | 2 | 33.57 | <0.001 |
| group × concentration | 3 | 46.63 | <0.001 |
| group × HM | 6 | 12.41 | 0.053 |
| concentration × HM | 2 | 73.82 | <0.001 |
| group × concentration × HM | 6 | 19.4 | 0.0035 |

Biomass was analysed using linear mixed-effects models fitted on both the full dataset and on a subset including only surviving individuals. Patterns were generally consistent between the two datasets (Table 2). In both cases, likelihood ratio tests supported the inclusion of the three-way interaction between group, metal, and concentration (all data: χ² = 35.36, df = 6, *p* < 0.001; survival-only: χ² = 19.85, df = 6, *p* = 0.003), indicating that the effect of concentration on biomass varied among groups and metals.

To facilitate interpretation of the three-way interaction, responses were examined separately for each metal (Table S4; Figure 3). In copper, biomass decreased with increasing concentration in all groups, with a weaker decline in *F. oxysporum* isolates from the mining environment (Figure 3A). The group × concentration interaction was significant in the full dataset but not in the biomass dataset (Table S4). In iron, biomass decreased across all groups with limited differences between groups (Figure 3A), and the group × concentration interaction was not significant when restricting the analysis to surviving individuals (Table S4). In zinc, biomass responses differed among groups, with *F. oxysporum* mining isolates showed increased biomass at higher concentrations, whereas all other groups exhibited decreasing trends (Figure 3A). This pattern was consistent between biomass of all data and biomass of surviving isolates only, with a significant group × concentration interaction in each case (Table S3, Figure S1).

**Figure 3:**
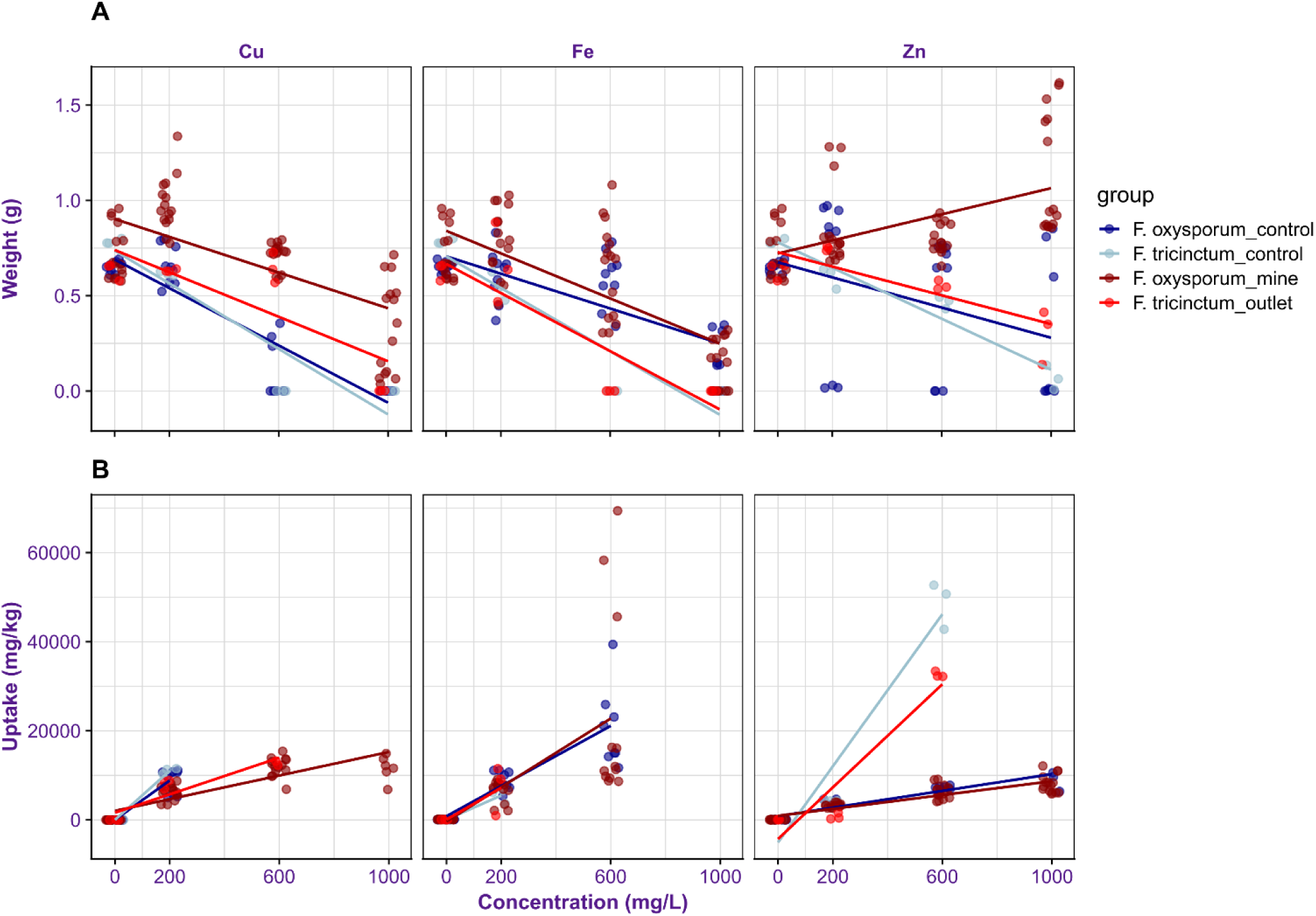
Overview of biomass and uptake for fungal isolates isolated from the mining environment and closely related controls. *Fusarium* isolates from the mine are shown in red, with closely related control isolates in blue. A) Biomass (dry weight in g) of all fungal isolates with > 0 g of dry weight. B) Relative uptake (mg metal per kg of dry weight mycelium) in fungal samples with > 3.5 g fresh weight.

### Reduced relative uptake of metals in several mine isolates

For all metals, *F. oxysporum* control isolates showed a higher relative uptake at 200mg/L than mine isolates (Figure 3B), this trend was also seen at higher zinc concentrations. Mine isolates showed a higher average relative iron uptake at 600 mg/L enrichment. This trend was driven by a single mine isolate, I3, with exceptionally high uptake. All other mine isolates had a lower relative uptake of iron than control isolates, suggesting that the apparent higher uptake in mine fungi at 600 mg/L iron may be due to variability among isolates rather than a general trend. Like *F. oxysporum*, the *F. tricinctum* outlet isolate maintained enhanced growth under metal stress while exhibiting reduced relative uptake of copper and zinc (Figure 3A and 3B). A higher average iron uptake was found in the *F. tricinctum* outlet isolate compared to the control isolate.

Both mine *F. oxysporum* isolates and the outlet *F. tricinctum* isolate showed a lower relative uptake of metals in comparison to control isolates at comparable concentrations, even in MEB media. Although several isolates from metal-enriched environments took up less heavy metal on average, general uptake levels were highly isolate specific, highlighting the importance to assess individual isolates to evaluate mycoremediation potential.

### Absolute metal uptake is largely isolate dependent

Absolute metal uptake appeared to be isolate-dependent and varied according to environmental origin, metal identity and concentration (Table 3). Differences between *F. oxysporum* control and mine isolates were negligible at low concentration (200 mg) for iron and copper, but control isolates took up slightly more zinc on average. For iron, the mine isolate I3 and the control isolate CBS149.25 showed exceptional iron uptake at intermediate levels (600 mg/L iron, accumulation of 23 mg and 16 mg, respectively), while the control isolate CBS151.27 showed relatively high amounts of iron uptake across a range of concentrations, with an average uptake of 9-10 mg. Mine isolate I4 demonstrated the highest copper uptake, maintaining robust growth up to 1000 mg/L and accumulating 7–10 mg across all tested concentrations. All *F. oxysporum* mine isolates grew well at high levels of zinc. Mine isolate I6 showed a generally large uptake across all levels of zinc enrichment, ranging from 4 mg in 200 mg/L enrichment to 11 mg in 1000 mg/L enrichment (Table 3). Some *F. oxysporum* isolates showed notably lower levels of absolute uptake than others. For example, mine isolate I7 showed low average iron uptake (1.8 mg) compared to all other *F. oxysporum* isolates (range from 4.4 – 9.3 mg uptake) at 200 mg/L iron enrichment

**Table 3:** Absolute metal uptake in fungal samples showing ≥ 3.5 g fresh weight.

| Absolute uptake (mg) |  |  |  |  |  |  |  |
| --- | --- | --- | --- | --- | --- | --- | --- |
| Metal | Species | Group | isolate | MEB | 200 mg/L | 600 mg/L | 1000 mg/L |
| Iron | *F. oxysporum* | Control | CBS 106.17 | 0.051 ± 0.00 | 6.734 ± 0.47 | 11.739 ± 8.82 |  |
|  |  | Control | CBS 149.25 | 0.068 ± 0.00 | 4.390 ± 1.27 | 16.052 ± 1.54 |  |
|  |  | Control | CBS 151.27 | 0.042 ± 0.00 | 9.229 ± NA | 9.570 ± 0.47 |  |
|  |  | Mine | I3 | 0.065 ± 0.00 | 4.582 ± 1.02 | 23.442 ± 1.03 |  |
|  |  | Mine | I4 | 0.069 ± 0.00 | 5.498 ± 0.54 | 12.052 ± 3.22 |  |
|  |  | Mine | I5 | 0.068 ± 0.00 | 7.537 ± 0.60 |  |  |
|  |  | Mine | I6 | 0.053 ± 0.01 | 7.662 ± 1.26 | 7.232 ± 0.78 |  |
|  |  | Mine | I7 | 0.052 ± 0.00 | 1.824 ± 0.92 | 8.860 ± 2.61 |  |
|  | *F. tricinctum* | Control | CBS 393.93 | 0.040 ± 0.00 | 2.924 ± 0.58 |  |  |
|  |  | outlet | I8 | 0.043 ± 0.00 | 5.484 ± 4.88 |  |  |
| Copper | *F. oxysporum* | Control | CBS 106.17 | 0.005 ± 0.00 | 6.691 ± 0.45 |  |  |
|  |  | Control | CBS 149.25 | 0.005 ± 0.00 | 2.946 ± 0.34 |  |  |
|  |  | Control | CBS 151.27 | 0.004 ± 0.00 | 8.485 ± 0.07 |  |  |
|  |  | Mine | I3 | 0.003 ± 0.00 | 6.508 ± 0.33 | 8.835 ± 0.54 | 5.537 ± 0.50 |
|  |  | Mine | I4 | 0.004 ± 0.00 | 7.426 ± 0.36 | 9.946 ± 0.41 | 9.173 ± 1.38 |
|  |  | Mine | I5 | 0.004 ± 0.00 | 7.286 ± 0.29 | 6.944 ± 1.29 |  |
|  |  | Mine | I6 | 0.003 ± 0.00 | 3.941 ± 0.38 | 8.687 ± 0.59 |  |
|  |  | Mine | I7 | 0.003 ± 0.00 | 5.158 ± 0.65 | 8.625 ± 0.53 |  |
|  | *F. tricinctum* | Control | CBS 393.93 | 0.003 ± 0.00 | 6.799 ± 0.44 |  |  |
|  |  | outlet | I8 | 0.003 ± 0.00 | 5.154 ± 0.67 | 8.730 ± 1.25 |  |
| Zinc | *F. oxysporum* | Control | CBS 106.17 | 0.035 ± 0.00 | 2.848 ± 0.19 | 5.503 ± 0.41 | 6.554 ± 1.22 |
|  |  | Control | CBS 149.25 | 0.036 ± 0.00 |  |  |  |
|  |  | Control | CBS 151.27 | 0.036 ± 0.00 | 4.209 ± 0.45 | 4.891 ± 0.20 |  |
|  |  | Mine | I3 | 0.034 ± 0.00 | 2.255 ± 0.03 | 4.193 ± 0.30 | 9.586 ± 2.38 |
|  |  | Mine | I4 | 0.034 ± 0.00 | 2.970 ± 0.13 | 5.505 ± 0.24 | 5.895 ± 1.17 |
|  |  | Mine | I5 | 0.039 ± 0.00 | 2.531 ± 0.10 | 5.506 ± 0.31 | 10.850 ± 1.64 |
|  |  | Mine | I6 | 0.032 ± 0.00 | 3.853 ± 0.10 | 7.112 ± 1.69 | 10.492 ± 1.70 |
|  |  | Mine | I7 | 0.036 ± 0.00 | 1.933 ± 0.02 | 3.122 ± 0.38 | 7.921 ± 1.80 |
|  | *F. tricinctum* | Control | CBS 393.93 | 0.039 ± 0.00 | 2.600 ± 0.29 | 22.806 ± 3.77 |  |
|  |  | outlet | I8 | 0.039 ± 0.00 | 0.568 ± 0.57 | 18.073 ± 0.62 |  |

The *F. tricinctum* control isolate tended to accumulate more copper and zinc at low levels of supplementation (200 mg/L), while the outlet isolate took up relatively more iron. Both *F*. *tricinctum* isolates showed a unique potential for zinc uptake at medium enrichment levels; control species took up 23 mg of zinc on average, compared to 18 mg for the outlet isolate at a concentration of 600 mg/L (Table 3).

## Discussion

Previous studies have shown that fungal populations from contaminated environments often show high metal tolerance through physiological and/or genetic adaptations (13,34), but direct comparison between isolates from mining environments and closely related controls have remained limited to date. In this study, we obtained six *Fusarium* isolates from sediments inside, and at the outlet of, the Kiirunavaara iron ore mine in Northern Sweden. Most heavy metals are enriched in the mine compared to the reference lake, the Norrbotten county average, and the national average. For some metals, the mine and outlet display lower levels of heavy metals (29). As the average for Norrbotten county and the national average contain a wide range of samples from various environments, these higher heavy metal levels are most likely the result of leaching from mines, roads, or other industrial runoff. Most importantly, the Kiirunavaara mine showed elevated concentrations of heavy metals in comparison to its outlet, indicating the potential for fungal populations inside the mine to adapt to higher levels of heavy metals than those at the outlet or in non-contaminated environments. By comparing isolates from the mining environment with closely related control isolates, we investigated if environmental origin was associated with differences in fungal performances under heavy metal exposure.

Fungal isolates from the mining environment and from the mine outlet showed increased survival and biomass compared to control isolates, indicating greater tolerance to heavy metals. Toxicity levels were furthermore metal dependent (35). The toxicity levels followed the order Cu > Fe > Zn, suggesting the presence of metal-specific tolerance mechanisms. The observed patterns are consistent with previous observations that environmental exposure can select for metal-resistant phenotypes. Although a minimum inhibitory concentration of 100 mg/L for zinc has been reported for one *F. oxysporum* isolate (35), the *F. oxysporum* isolates from the Kiirunavaara mine exhibited substantially higher tolerance, maintaining elevated dry weight even at the highest tested concentration (1000 mg/L zinc). Similarly, the *F. tricinctum* isolate from the outlet also displayed greater metal tolerance than its corresponding control, although to slightly lower levels than *F. oxysporum*.

We observed inherent taxon-level differences in metal sensitivity. The *F. tricinctum* isolate from the mine outlet, where metal concentrations are lower than in the mine pumping pits, exhibited lower biomass at higher metal concentrations than the *F. oxysporum* isolates from the mine. Although species identity and environmental origin were confounded in this dataset, *F. oxysporum* generally exhibited greater metal tolerance than *F. tricinctum*, including when control isolates were compared directly. These findings are consistent with previous studies reporting species-specific differences in metal resistance among fungi inhabiting similar environments (14,36). Considering the difference in metal tolerance amongst different isolates, future studies could test tolerance to multiple heavy metals. Such a multi-metal analysis would more closely reflect in-situ conditions of the mining environment, where fungi are exposed to complex mixtures of cobalt, nickel, vanadium and other metals. In addition, it would be of interest to assess growth in-situ and control for pH. In this experiment, the addition of the metal solutions lowered the pH of the media, with the lowest pH (pH 3.35) found in 1000 mg/L copper treatment. While *Fusaria* and other filamentous fungi have been shown to be able to grow in acidic conditions (37,38), pH also affects metal toxicity (39). Thus, the inhibitory effects observed here could reflect the combined effects of metal concentration and pH. Future investigations in which pH is adjusted independently of metal concentration would be helpful to disentangle the effects of pH from those of heavy metal toxicity.

This comparison between isolates from within and outside the mine are consistent with the hypothesis that fungal tolerance increases in environments subjected to greater metal-related selective pressure. However, the limited number of independent isolates included in this study, particularly for *F. tricinctum*, does not allow for definitive conclusions regarding local adaptation. In addition, since fungal identity and environmental origin were not fully crossed in our experimental design, it is not possible to disentangle the relative contributions of these factors. Thus, the differences should be interpreted as associations between environmental origin and fungal performance, rather than direct evidence of adaptive evolution.

### Relationship between heavy metal tolerance and heavy metal uptake

In congruence with earlier studies by Smith et al., 2024 (16), isolates from the mine and outlet showed lower relative uptake levels than control isolates, potentially reflecting adaptive strategies for surviving in toxic environments (15). To this end, fungi possess dedicated homeostatic systems to control import, export, storage and transport of metal ions (40). Under long-term exposure to contaminated environments, these homeostatic mechanisms can evolve to reduce uptake and enhance detoxification mechanisms, allowing survival under otherwise toxic conditions (40,41). For example, copper tolerance in *Fusarium* species has been linked to Cu^+^ efflux systems and activation of oxidative stress responses (42,43). Zinc resistance is often associated with increased efflux of zinc ions, vacuolar storage, and zinc chelation that immobilizes excess zinc (17,44). Additionally, although data on iron resistance in filamentous fungi remains limited, studies in yeast suggest similar mechanisms that involve reduced uptake and transport to the vacuoles (40). Indeed, yeasts such as S*accharomyces cerevisae* have been reported to have an inverse relationship between uptake and survival in heavy metal enrichment, most likely caused the lack efflux pathways and/or the lack of pathways to exclude metals from the cell in the more sensitive strains (45).

Further research combining physiological assays with gene expression analyses would help clarify whether performance in these isolates arises mainly from reduced uptake, enhanced efflux, intracellular detoxification, or a combination of these pathways. Such studies would further help determine whether the observed differences in survival, biomass, and metal uptake amongst control and mine isolates are primarily driven by environmental exposure to heavy metals or by substrate origin. This study compared *Fusarium* isolates from mine sediments with closely related isolates obtained from plants, and in one case obtained from an unknown origin. Some physiological pathways involved in heavy metal tolerance of *F. oxysporum*, such as copper reductase and copper transporters, may be active specifically during plant infection (46). Since all the isolates were cultured under identical laboratory conditions, we assume that differences related to the original substrate (i.e., plants or other environments) have been minimized and expect the observed differences to reflect intrinsic differences in performance under stress from heavy metals.

### Absolute uptake is isolate dependent

While previous studies have highlighted filamentous fungi from polluted environments as promising candidates for mycoremediation, our findings indicate that mycoremediation potential, which we assessed in this study as absolute uptake of a metal in the grown mycelium, is not necessarily linked to environmental origin. Instead, survival, biomass, and relative uptake varied substantially amongst isolates and species. The mine isolates that survive high levels of metal enrichment may not always be the best myco-accumulators, and some control isolates very effectively removed metals under moderate enrichment. Additionally, some isolates show exceptional uptake at a specific level of enrichment, while others show a relatively large uptake across a range of enrichment levels.

As such, different isolates may serve different purposes, depending on the origin and level of contamination. The high uptake of control isolates might serve a unique purpose in quickly eliminating heavy metals from moderately contaminated environments, while the survival and growth of mine-derived fungi could serve as valuable bioresources for detoxifying contaminated sites, particularly in scenarios where survival under extreme heavy metal stress is required. Research should be conducted into specific isolates to further assess mycoremediation capabilities. Specifically, *F. oxysporum* isolates CBS 151.27, I4, and I6 should be further researched for iron, copper and zinc uptake, respectively, over a span of different heavy metal concentrations. At the same time, I3 should be further researched for iron uptake, specifically at levels of 600 mg/L enrichment. Lastly, the *F. tricinctum* control isolate CBS 393.93 could provide a unique opportunity for mycoremediation at zinc concentrations of 600 mg/L. Genomic and transcriptomic studies in these isolates could help to identify the mechanisms underlying the relationship between heavy metal tolerance and accumulation, providing further ecological insights and driving practical bioremediation strategies. Together, these approaches will enhance the development of fungal-based strategies for effective heavy metal detoxification across diverse contamination scenarios.

In this study, fungal cultures were mechanically cleaned and assumed to be free of bacterial contaminants based on absence of bacterial PCR products before onset of the experiment. This allows for analyses of mycoremediation potential by fungal monocultures. However, fungi in mining environments naturally occur within complex microbial communities which include both fungal and bacterial taxa (47), where interspecific interactions can influence both metal tolerance and uptake. Increasing evidence suggests that co-cultures of fungi and bacterial partners may outperform monocultures (48). Such community-based approaches represent a promising direction to reveal complementary bioremediation strategies (40,41).

## Conclusion

This study demonstrates that fungal isolates from contaminated mining environments exhibit enhanced tolerance to heavy metals, as reflected by increased survival and biomass under elevated concentrations of copper, iron, and zinc (up to 1000 mg/L enrichment). For both species, differences in survival and biomass show a toxicity gradient Zn < Cu < Fe. Among the tested isolates, *Fusarium oxysporum* from the mine showed the highest survival, with strong resistance to copper and even higher biomass under high zinc concentrations. Tolerant isolates from the mining environment displayed lower uptake of heavy metals into the mycelium. We thus conclude that suitability for mycoremediation potential should be evaluated on a case-by-case basis for fungal isolates, considering both tolerance and relative uptake rather than environmental origin alone.

## Acknowledgements

Ethics approval and consent to participate

Not applicable.

We thank Anbar Khodabandeh for assistance in fungal isolation and laboratory experiments.

## Consent for publication

Not applicable.

## Availability of data and materials

Raw data is available in table S2.

## Author contributions

PM and NH designed the study and its experiments. All authors contributed to the study design and conceptualization. PM and NH performed sediment collection and laboratory work, and were main contributors in statistical analysis, data interpretation and writing of the manuscript. PM performed fungal isolations. MO performed statistical analysis and data interpretation. All authors contributed to text-editing and approved the final manuscript.

