## supplementary methods for "Living in a metal-rich world: Enhanced growth and reduced metal accumulation in *Fusarium* fungi from the Kiirunavaara iron ore mine"

### Supplementary methods for DNA extraction and PCR analysis of fungal isolates from the Kiruna iron ore mine

For DNA extraction, a small amount of mycelium from each fungal culture was added separately to 50-100 µL of 10% Chelex solution in 96-well plates. Negative controls were prepared without the addition of mycelium. The plates were vortexed for 10-20 seconds, centrifuged briefly, and incubated at 95°C for 30 minutes. The resulting DNA extracts were used for subsequent PCR analysis of the internal transcriber spacer (ITS) region for fungal identification, and the 16S region to check for bacterial contamination, as described by Klindworth et al. (2013). PCR reactions were set up using 1:1 diluted DNA extract. For reactions where no product was initially obtained, undiluted samples were tested. A PCR master mix (50 µL) was created with 10 µL 5x Phusion HF Buffer, 1 µL 10 mM dNTP, 1.5 µL DMSO, 0.5 µL Phusion DNA Polymerase, and 0.5 µL of each of the two primers. The PCR reaction was performed on 49 µL of master mix and 1 µL of DNA template (1).

The amplification of the ITS region was performed with primers ITS1 and ITS4 (Supplementary table 3), and the following PCR cycle: Initial denaturation at 98°C for 1 minute, 35 cycles of denaturation at 98°C for 15 seconds, annealing at 58°C for 30 seconds, and extension at 72°C for 45 seconds, followed by a final extension at 72°C for 7 minutes. For cleaning, fungal ITS PCR products were diluted 1:8, and 10 µL of the diluted PCR product was mixed with 3 µl of ExoProStar reagent. The mixture was incubated at 37°C for 30 minutes, followed by 80°C for 15 minutes. Five µL of the purified PCR product was transferred into a 1.5 mL tube and prepared for sequencing using 5 µM primer. ITS sequences of the pure cultures of the original fourteen morphotypes were sequenced with Sanger sequencing (Eurofins). Species identity was inferred by comparing obtained ITS sequences to the NCBI database using BLASTN (2).

Potential bacterial contamination was identified by testing for presence of 16S rDNA with two primer-pairs, the primer BC1 pair S-D-Bact-0341-b-S-17 and S-D-Bact-0785-a-A-21, and the primer BC4 pair SD-Bact-0008-a-s-16 and SD-Bact-0907-a-A-20, as described by Klindworth et al.

(2013)(Table 1). The BC1 PCR program used initial denaturation at 98°C for 1 minute, followed by 35 cycles of denaturation at 98°C for 30 seconds, annealing at 68°C for 30 seconds, and extension at 72°C for 30 seconds, with a final extension at 72°C for 7 minutes. The BC4 program used initial denaturation at 98°C for 1 minute. 35 cycles of denaturation at 98°C for 15 seconds, annealing at 52°C for 30 seconds, and extension at 72°C for 30 seconds, with a final extension at 72°C for 7 minutes. Gel electrophoresis was used to confirm absence of bacterial contaminants. Fungal cultures were assumed to be pure, based on absence of bacterial PCR products at the start of the experiment.

Table 1: Primers used for ITS-sequencing of fungal isolates and 16S rDNA primers used to identify bacterial DNA .

| | | | |
|---|---|---|---|
| ITS primers | ITS1 | | (5'-TCCGTAGGTGAACCTGCGG-3') |
| | ITS-4 | | (5'-TCCTCCGCTTATTGATATGC-3') |
| 16S primers | BC1 pair | S-D-Bact-0341-b-S-17 | (5'-CCTACGGGNGGCWGCAG-3') |
| | | S-D-Bact-0785-a-A-21 | (5'-GACTACHVGGGTATCTAATCC-3') |
| | BC4 pair | SD-Bact-0008-a-s-16 | (5'-AGAGTTTGATCMTGGCTCAG-3') |
| | | SD-Bact-0907-a-A-20 | (5'-CCGTCAATTCMTTTGAGTTT-3') |
