## Supplementary material for "Living in a metal-rich world: Enhanced growth and reduced metal accumulation in *Fusarium* fungi from the Kiirunavaara iron ore mine": Figure S1

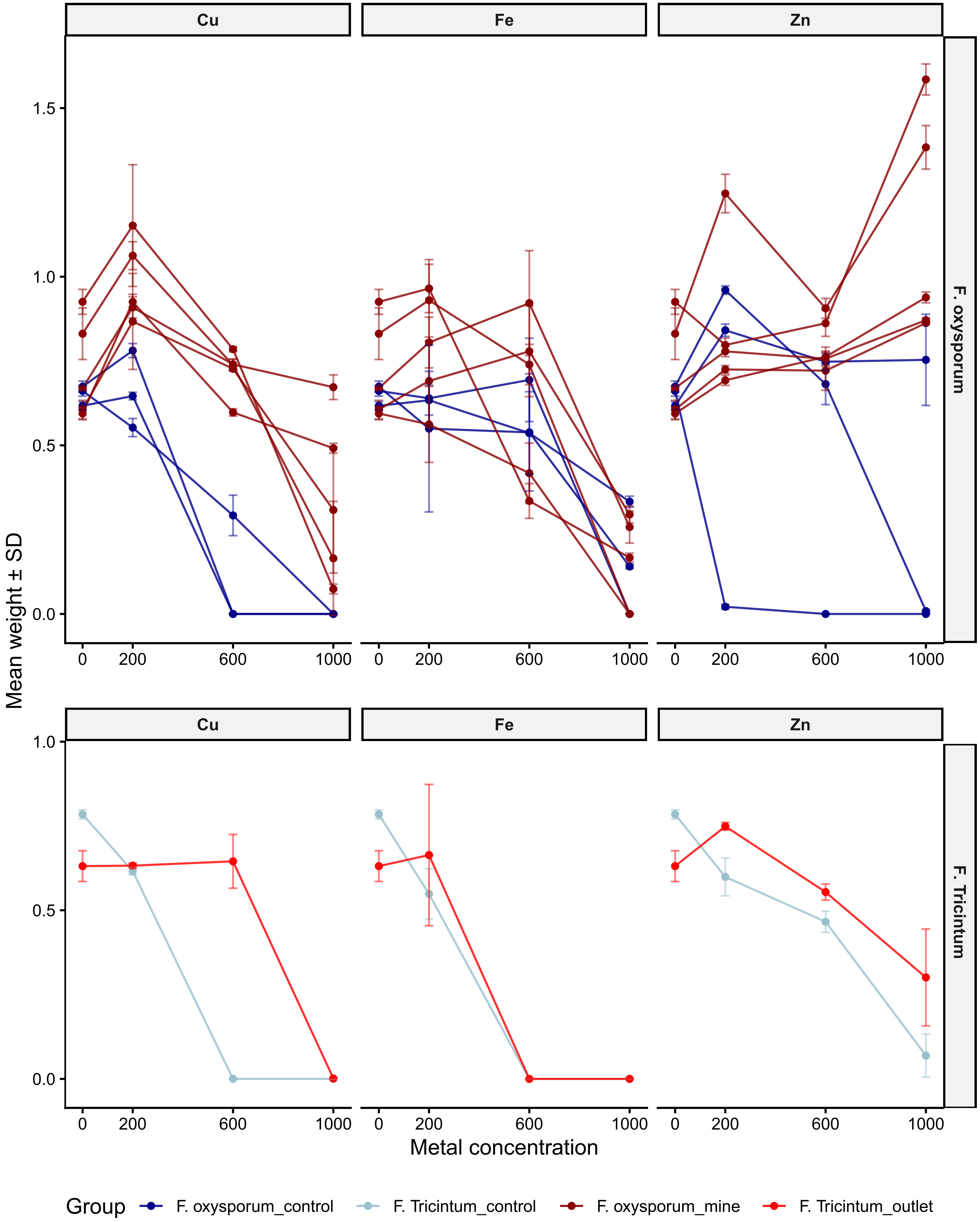


Figure S1: dry weight +- standard deviation (in grams) of assessed strains for *Fusarium oxyporum* (upper panel) and *Fusarium tricintum* (lower panel). Individual strains are shown as individual lines, with standard deviation based on the three assessed replicates. Dry weights of individual strains, including strain identification, are shown in Table S3.
